# AT-hook-dependent DNA loop extrusion by STAG1 drives 3D genome folding

**DOI:** 10.1101/2025.11.17.688764

**Authors:** Ryota Sakata, Daniel Giménez, Ana Losada, Tomoko Nishiyama

## Abstract

Cohesin, a ring-shaped complex composed of Smc1, Smc3, Scc1 and STAG, is essential for sister chromatid cohesion and the regulation of three-dimensional (3D) genome architecture. At the single-molecule level, cohesin extrudes DNA loops, a process thought to drive higher-order genome folding. In vertebrates, cohesin incorporates either STAG1 or STAG2. Although both support sister chromatid cohesion, they differentially regulate 3D genome organization. However, the mechanistic basis for these differences has remained unclear. Here we show, using single-molecule assays, that cohesin-STAG1 extrudes DNA loops more efficiently than cohesin-STAG2, despite comparable ATPase activity and topological DNA entrapment. We identify an AT-hook motif unique to the STAG1 N-terminus as the element that promotes loop extrusion without altering ATPase activity or DNA binding. In human somatic cells, the AT-hook is required for stable cohesin-chromatin association during G1 phase but is dispensable for sister chromatid cohesion. Mutation of this motif markedly impairs TAD and chromatin loop formation. These findings highlight AT-hook as a critical determinant that distinguishes STAG1 from STAG2 by promoting DNA loop extrusion and stabilizing cohesin-chromatin interactions in interphase through a mechanism distinct from the one underlying sister chromatid cohesion.

## Introduction

Cohesin, a ring-shaped chromosomal ATPase that consists of four core subunits (Smc1, Smc3, Scc1, and stromal antigen (STAG)), topologically entraps DNA to achieve sister chromatid cohesion^1,2^. Recently, numerous studies have highlighted that cohesin plays essential roles in building up higher-order chromatin structures such as topologically associating domains (TADs), A/B compartments and chromatin loops^3,4^. DNA loop extrusion has been proposed as putative mechanism for this function^5,6^, yet direct evidence for how this activity contributes to genome organization is still lacking.

Only limited information is so far available on the mechanistic similarities and differences between sister chromatid cohesion and DNA loop extrusion. From a structural perspective, it has been established that all four cohesin core subunits as well as the ATPase activity within the Smc1/3 head domains are required for both processes^5–8^. On the other hand, topological entrapment of DNA by cohesin is essential only for cohesion^7^ but not for loop extrusion^5^. In addition, a DNA-binding patch formed by the non-SMC subunits Scc1 and STAG is necessary for interactions between cohesin and DNA, as well as with various cohesin regulators such as CCCTC-binding factor (CTCF), Sgo1, and Wapl^9–13^. This DNA-binding patch is required for DNA loop extrusion^14,15^ but not for topological loading^15^ . Conversely, a basic patch within the Smc1/3 hinge domain is essential for cohesion but not for loop extrusion^16^. Furthermore, biochemical analyses and single-molecule imaging suggest that loop extrusion relies on flexible and dynamic structural rearrangements of the cohesin complex^14,17,18^ . Taken together, while multiple lines of evidence point to cohesion and loop extrusion being mechanistically distinct, the precise molecular basis of these differences remains unresolved.

In vertebrate, cohesin complex possesses one of two mutually exclusive HEAT repeat subunits, STAG1 or STAG2^19,20^ (Fig.1a), while germ cells carry also STAG3^21^. STAG1 and STAG2 are encoded by paralogous genes and are structurally similar, except for their intrinsically disordered regions (IDR) at the N- and C-terminus, but several studies have highlighted their functional divergence^22^. While both STAG1 and STAG2 can establish sister chromatid cohesion^23^, they have more prominent roles in telomere and centromere cohesion, respectively ^24–26^. Loss of STAG1 alters long-range genome interaction, weakens TADs boundaries and increases compartmentalization^27–29^, whereas STAG2 loss primarily disrupts local contacts and affects cell type-specific gene regulation^29–31^, indicating that they contribute to genome organization distinctly. Furthermore, cohesin-STAG1 associates more stably with chromatin than cohesin-STAG2^27,28^, possibly because cohesin-STAG1 is preferentially located at CTCF-binding sites and acetylated by Esco1 than cohesin–STAG2^27,28,32^. Although accumulating evidence highlights their distinct modes of regulation in higher-order chromatin structure, the molecular determinants underlying these differences remain unclear.

Here, we investigate how STAG1 and STAG2 differentially regulate the molecular activity of cohesin. Single-molecule assays reveal that cohesin-STAG1 extrudes DNA loops more efficiently and binds DNA more stably than cohesin-STAG2, despite comparable ATPase activity and topological DNA binding. We identify the STAG1-specific AT-hook motif as a key enhancer of loop extrusion and post-loading stabilization. In cells, AT-hook promotes stable chromatin binding of cohesin-STAG1 during G1 phase by facilitating Smc3 acetylation and CTCF interaction, whereas in G2 phase, stabilization by Sororin renders AT-hook dispensable for cohesion. AT-hook mutation impairs 3D genome organization, weakening TADs and chromatin loops. Together, these findings uncover a mechanistic basis for the functional divergence of STAG1 and STAG2, establishing paralog-specific loop extrusion as a central driver of genome organization.

## Results

### Cohesin-STAG1 and cohesin-STAG2 exhibit equivalent topological DNA binding activity

To investigate the mechanistic differences between STAG1 and STAG2, we examined their topological DNA binding ability, which is essential for sister chromatid cohesion^33^, using an *in vitro* assay previously described^7^. Recombinant human trimeric cohesin complex lacking a STAG protein, tetrameric cohesin containing either STAG1 or STAG2 (hereafter referred to as cohesin-STAG1 or cohesin-STAG2; Fig. 1a,b), and NIPBL-Mau2, the cohesin loader, were purified from insect cells (Fig.1c). Next, trimeric or tetrameric cohesin was incubated with relaxed circular DNA in the presence of NIPBL-Mau2 and ATP. After the high-salt wash to remove unbound DNA, cohesin was immunoprecipitated and the amount of retained DNA was quantified (Fig. 1d). Consistent with a previous study in yeast^7^, tetrameric cohesin successfully co-immunoprecipitated DNA even after the high-salt wash, whereas trimeric cohesin failed (Fig. 1d). Moreover, the two tetramers (cohesin-STAG1 and cohesin-STAG2) recovered DNA to a similar extent (Fig. 1d). BamHI digestion confirmed that the majority of bound DNA was topologically entrapped by cohesin-STAG1 and -STAG2 (Fig. 1e,f). These results demonstrate that the essential role of the STAG subunit for topological DNA binding, and that this function can be similarly performed by STAG1 and STAG2.

**Fig. 1.**
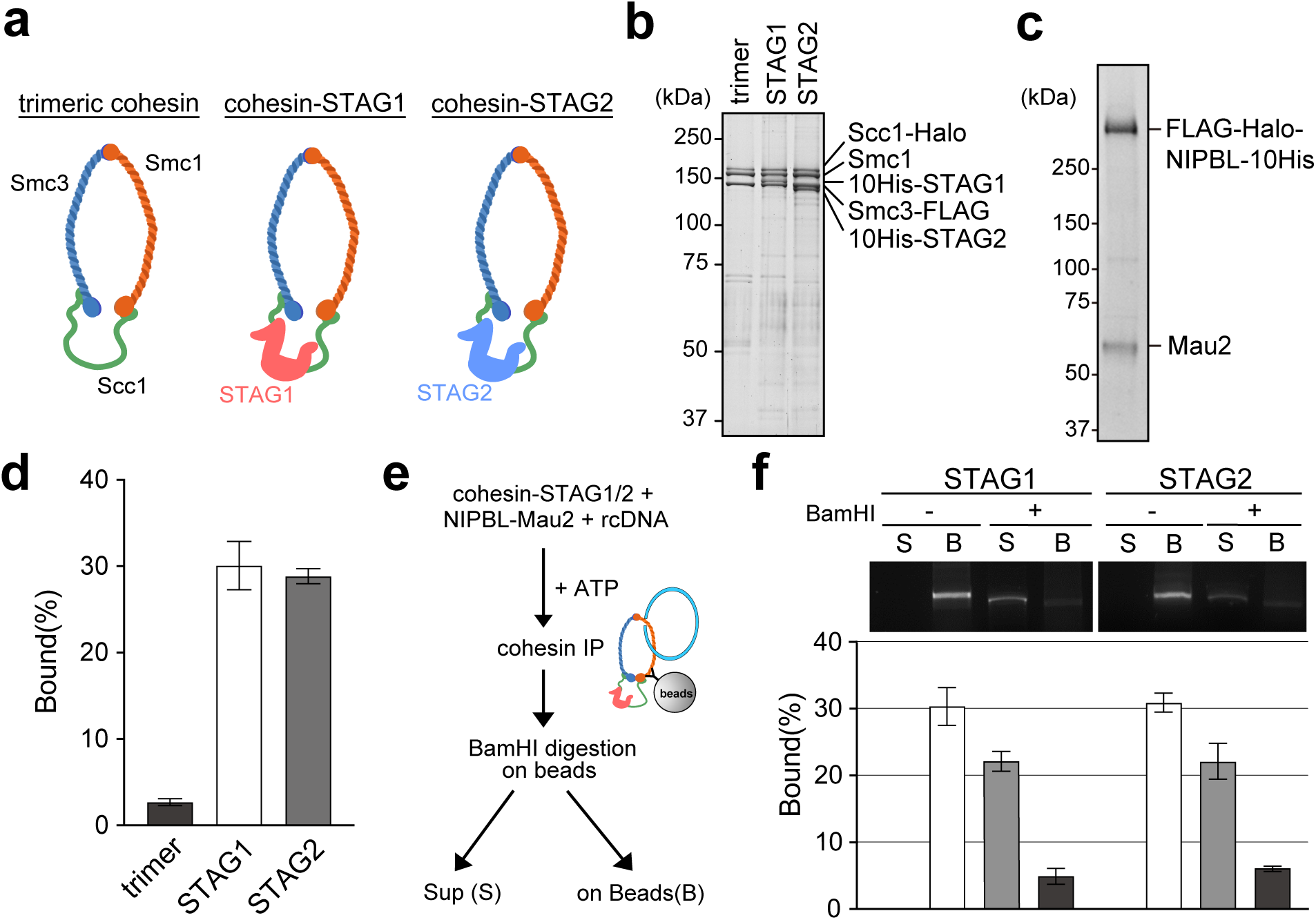
Cohesin-STAG1 and cohsin-STAG2 exhibit equivalent topological DNA binding activity. **a,** Schematic representation of cohesin complexes. **b**, **c,** SYPRO Ruby staining of the purified cohesin trimer, cohesin-STAG1, cohesin-STAG2 (**b**) and NIPBL-Mau2 complex (**c**) after separation by SDS-PAGE. **d,** *In vitro* topological DNA binding assay. Cohesin trimer or tetramer with STAG1 or STAG2 was loaded onto relaxed circular DNA (rcDNA) in the presence of NIPBL-Mau2 and ATP. Cohesin-bound DNA was retrieved with an antibody against Smc3, washed in high-salt buffer and subjected to electrophoresis, DNA intensities were quantified in 3 independent trials (mean ± SD). **e,f,** Schematic representation of the topological DNA binding assay followed by BamHI digestion (**e**). After cohesin immunoprecipitation (IP), cohesin-bound DNA was digested with BamHI to separate supernatant (S) and beads-bound (B) fractions. Quantification from 3 independent trials (mean ± SD) (**f**).

### Cohesin-STAG1 is a better DNA loop extruder than cohesin-STAG2

We next compared DNA loop extrusion activity of the two complexes. Although previous studies have implied that both cohesin-STAG1 and -STAG2 are capable of loop extrusion^5^, no direct molecular evidence has been provided so far. We performed single-molecule imaging as previously described^34^, using biotinylated λ-DNA tethered to an avidin-coated coverslip and extended into a U-shape by buffer flow in a microfluidic chamber. Purified cohesin trimer or tetramer harboring either STAG1 or STAG2 was introduced into the flow-cell along with NIPBL-Mau2 and ATP, and loop formation was monitored using total internal reflection fluorescence microscopy (TIRFM). (Fig. 2a,b). Consistent with a previous study^5^, cohesin tetramer, but not trimer, extrudes DNA loops (Fig. 2c), confirming the requirement of STAG for loop extrusion. Importantly, cohesin-STAG1 exhibited higher loop extrusion activity than cohesin-STAG2 (Fig. 2c). Quantitative analysis revealed that loop growth rate (Fig. 2d) and loop stability (Fig. 2e) were similar between the two complexes, suggesting that STAG1 facilitates loop extrusion in the initiation step rather than during subsequent loop growth or loop stabilization steps.

**Fig. 2.**
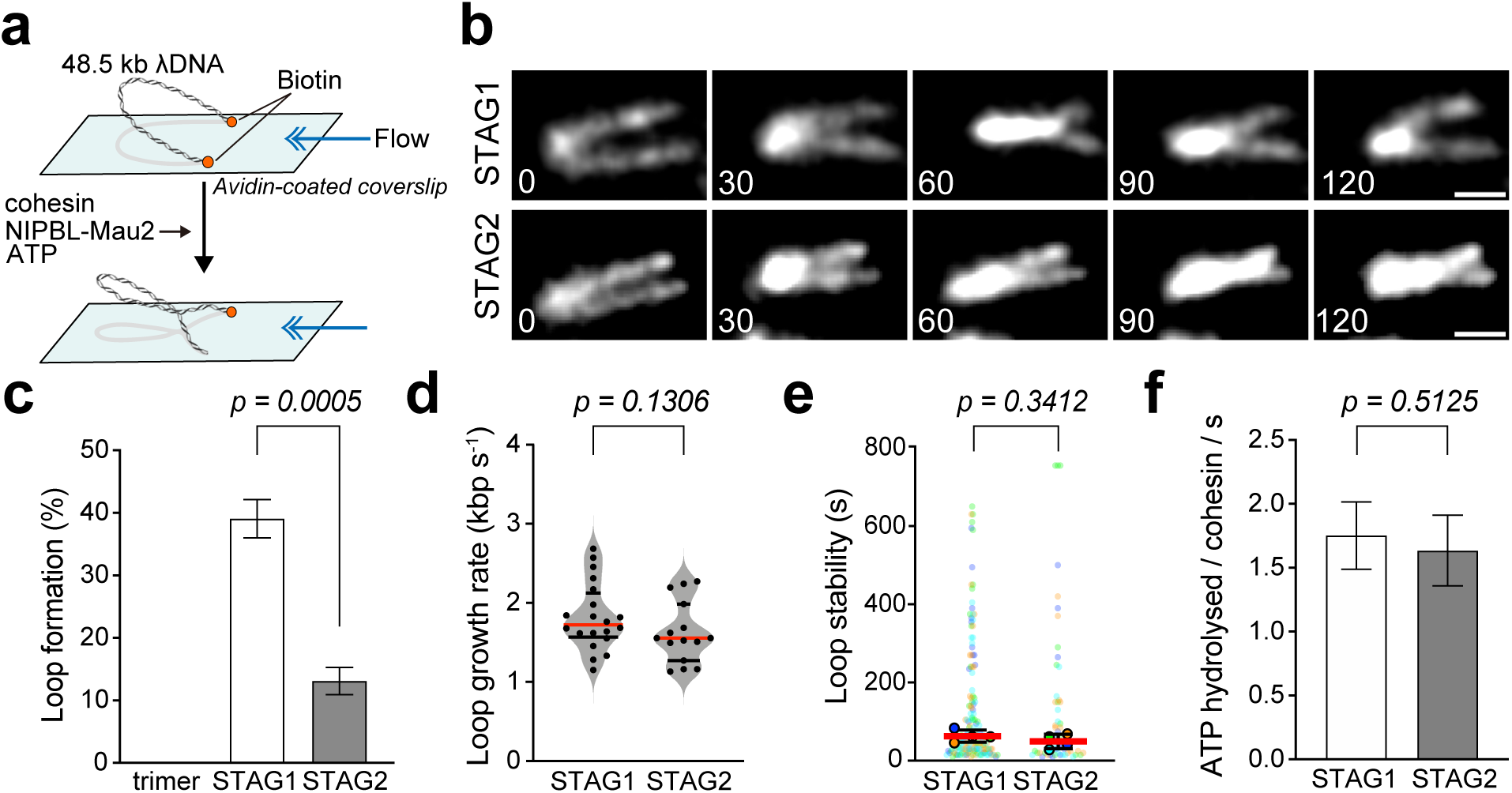
Cohesin-STAG1 is better DNA loop extruder than cohesin–STAG2. **a,** Schematic representation of DNA loop extrusion assay. Biotin-labeled λ-DNA molecules tethered on avidin-coated coverslip were stretched by perpendicular flow. DNA loop was formed in the presence of cohesin, NIPBL-Mau2 and ATP. **b,** DNA loop formed by cohesin-STAG1 (upper rows) and cohesin-STAG2 (lower rows). DNA was visualized by SYTOX Orange staining. Scale bar: 2 μm. **c,** Frequency of DNA loop formation on U-shaped DNA. Data were collected from 3 independent trials for trimer and 4 independent trials for tetramers (trimer : n = 105, 107, 117 DNAs; STAG1: n = 106, 96, 80, 90 DNAs; STAG2: n = 102, 99, 87, 76 DNAs, mean ± SD). The P value was assessed by Welch’s t test. **d,** DNA loop growth rate by cohesin-STAG1 or cohesin-STAG2. The red and black bars denote the median and quartiles, respectively. (STAG1: n = 20 DNAs; STAG2: n = 15 DNAs). The P values were assessed by a two-tailed Mann-Whitney U test. **e,** DNA loop stability assessed by DNA loop maintaining duration of each loop formed in the presence of cohesin-STAG1 or cohesin-STAG2. The solid point denotes the median of each condition. Corresponding data points are labeled with different colors for each experiment. The red and black bars denote the mean of the median and quartiles, respectively. Data were collected from 4 independent trials (STAG1: n = 28, 41, 52, 51 DNAs; STAG2: n = 14, 16, 17, 11 DNAs). The P values were assessed by a two-tailed Mann-Whitney U test. **f,** ATP hydrolysis rate of cohesin-STAG1 and cohesin-STAG2 in the presence of NIPBL-Mau2 and DNA. Data were collected from 3 independent trials (mean ± SD). The P value was assessed by Welch’s t test.

Loop extrusion strictly depends on cohesin ATPase activity^5,6^. We therefore tested whether the reduced loop extrusion activity of cohesin-STAG2 reflects lower ATPase activity. We found that both cohesin-STAG1 and -STAG2 exhibited a similar ATPase activity (Fig. 2f), which was stimulated by NIPBL-Mau2 and DNA (Extended Data Fig. 1). Thus, the lower loop extrusion activity of cohesin-STAG2 is not due to impaired ATPase activity. Together, these findings suggest that STAG1 promotes the initial step of loop extrusion more efficiently than cohesin-STAG2 by a mechanism that is independent of the ATPase activity.

### The AT-hook motif is crucial for promoting loop extrusion

Given the pronounced difference in loop extrusion activity between cohesin-STAG1 and -STAG2, we next sought to identify the molecular determinant underlying this difference. While STAG1 and STAG2 share over 75% sequence identity within their structured regions, their unstructured N- and C-terminal regions are far less conserved (less than 40% identity, Extended Data Fig. 2a). In the non-conserved N-terminal region of STAG1, there is an AT-hook DNA-binding motif (KRKRGRP)^35^, which associates with the DNA minor groove and is required for stable DNA binding^35–37^. To test if the AT-hook is required for STAG1-dependent efficient loop extrusion, we repeated the assay with tetrameric complexes harboring either STAG1 with a mutated AT-hook (STAG1^RA^) or STAG2 with an AT-hook inserted into the N-terminus (STAG2^AT^, Fig. 3a,b). Strikingly, loop formation efficiency was diminished in STAG1^RA^ and increased in STAG2^AT^ to levels closer to STAG1^WT^ (Fig. 3c). In contrast, neither loop growth rate nor loop stability was significantly affected (Fig. 3d,e). As expected, these mutations did not alter ATPase activity or its dependence on NIPBL-Mau2 and DNA (Fig. 3f, Extended Data Fig. 2b-d).

**Fig. 3.**
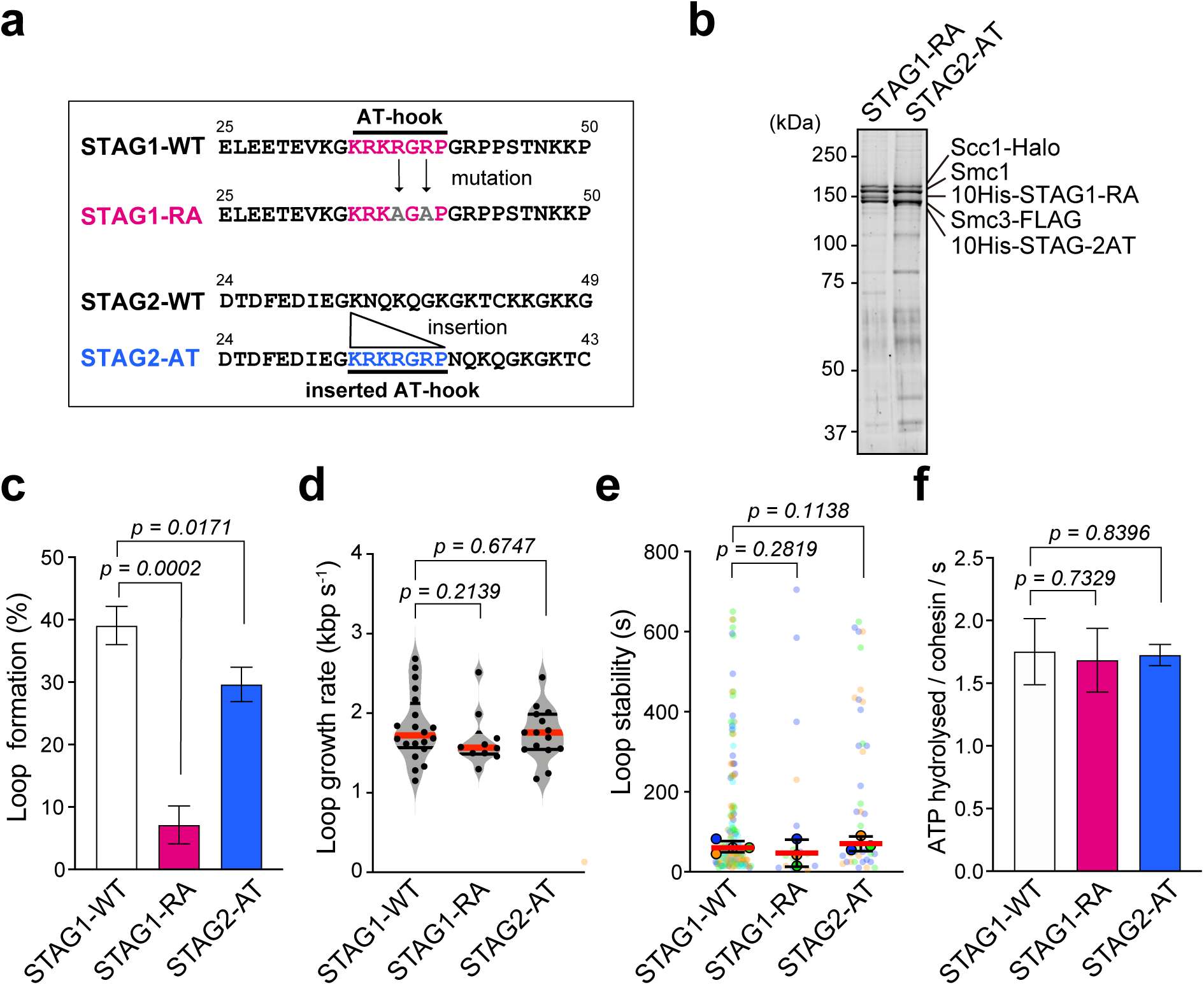
The AT-hook motif is crucial for promoting loop extrusion. **a,** Schematic representation of AT-hook mutated STAG1 (STAG1-RA) and AT-hook inserted STAG2 (STAG2-AT). In STAG1-RA, two arginine residues within the AT-hook motif were substituted with alanine. In STAG2-AT, the AT-hook sequence (KRKRGRP) was inserted into the corresponding N-terminal region of STAG2. **b,** SYPRO Ruby staining of the purified cohesin-STAG1-RA and cohesin-STAG2-AT after separation by SDS-PAGE. **c,** Frequency of loops on U-shaped DNA. Data of STAG1-WT, used for comparison, is same as **Fig.1c**. Data were collected from 3 independent trials (STAG1-RA: n = 76, 74, 76 DNAs; STAG2-AT: n = 65, 64, 36 DNAs, mean ± SD). The P value was assessed by Welch’s t test. **d,** DNA loop growth rate by cohesin-STAG1-WT, cohesin-STAG1-RA or cohesin-STAG2-AT. The red and black bars denote the median and quartiles, respectively. Data of STAG1-WT, used for comparison, is same as **Fig.1d**. (STAG1: n = 10 DNAs; STAG2: n = 15 DNAs). The P value was assessed by a two-tailed Mann-Whitney U test. **e,** DNA loop stability assessed by DNA loop maintaining duration of each loop formed in the presence of cohesin-STAG1-WT, cohesin-STAG1-RA or cohesin-STAG2-AT. The solid point denotes the median of each condition. Corresponding data points are labeled with different colors for each experiment. The red and black bars denote the mean of the median and quartiles, respectively. Data of STAG1-WT, used for comparison, is same as **Fig.1e**. Data were collected from 3 independent trials (STAG1-RA: n = 8, 4, 4 DNAs; STAG2-AT: n = 20, 14, 10 DNAs). The P value was assessed by a two-tailed Mann-Whitney U test. **f,** ATP hydrolysis rate of cohesin-STAG1-WT, cohesin-STAG1-RA and cohesin-STAG2-AT in the presence of NIPBL-Mau2 and DNA. Data of STAG1-WT, used for comparison, is same as **Fig.1f**. Data were collected from 3 independent trials (mean ± SD). The P value was assessed by Welch’s t test.

Next, we compared cohesin-STAG1 and -STAG2 dynamics on DNA at a single-molecule level. First, we evaluated DNA binding of fluorescently labeled cohesin on U-shaped λ-DNA in the presence of NIPBL-Mau2 and ATP (Extended Data Fig. 3a). DNA binding frequency was defined as the percentage of DNA molecules bound by cohesin, measured two minutes after the initial binding event. Unexpectedly, we observed no significant difference in binding frequency among STAG1^WT^, STAG2^WT^, STAG1^RA^ and STAG2^AT^ (Extended Data Fig. 3b). We then quantified the number of cohesin complexes bound to DNA by measuring fluorescence intensity (Extended Data Fig. 3c). Based on the average intensity of a single cohesin, determined by analyzing photobleaching steps of nonspecifically bound cohesin on the glass surface (Extended Data Fig. 3d,e), we calculated that one to two complexes bound per DNA molecule, on average, irrespective of paralogs or mutants (Extended Data Fig. 3c). These results indicate that AT-hook does not promote initial DNA binding of the cohesin. It is plausible that other cohesin subunits or NIPBL-Mau2 may compensate for AT-hook deficiency during initial DNA binding^38–^ _40._

### The AT-hook restricts cohesin dynamics on DNA

Since the AT-hook is not required for the initial DNA binding of cohesin, it may instead help stabilize cohesin after DNA binding. To test this possibility, we analyzed the diffusion of single cohesin molecules on stretched DNA. Cohesin-STAG1 or -STAG2 was bound to DNA in the presence of NIPBL-Mau2 and the non-specifically bound fraction was removed by high-salt wash. Subsequently, the diffusion of Halo-Alexa488-labeled cohesin was monitored by TIRFM (Fig. 4a). Both types of cohesin displayed diffusive motion as previously reported^41–43^ (Fig.4b). Notably, cohesin-STAG1 was more stably bound to DNA than -STAG2, as indicated by a lower diffusion coefficient (D = 0.1874 ± 0.0077 μm^2^/s for STAG1 vs. D = 0.2411 ± 0.0136 μm^2^/s for STAG2, Fig. 4c).

**Fig. 4.**
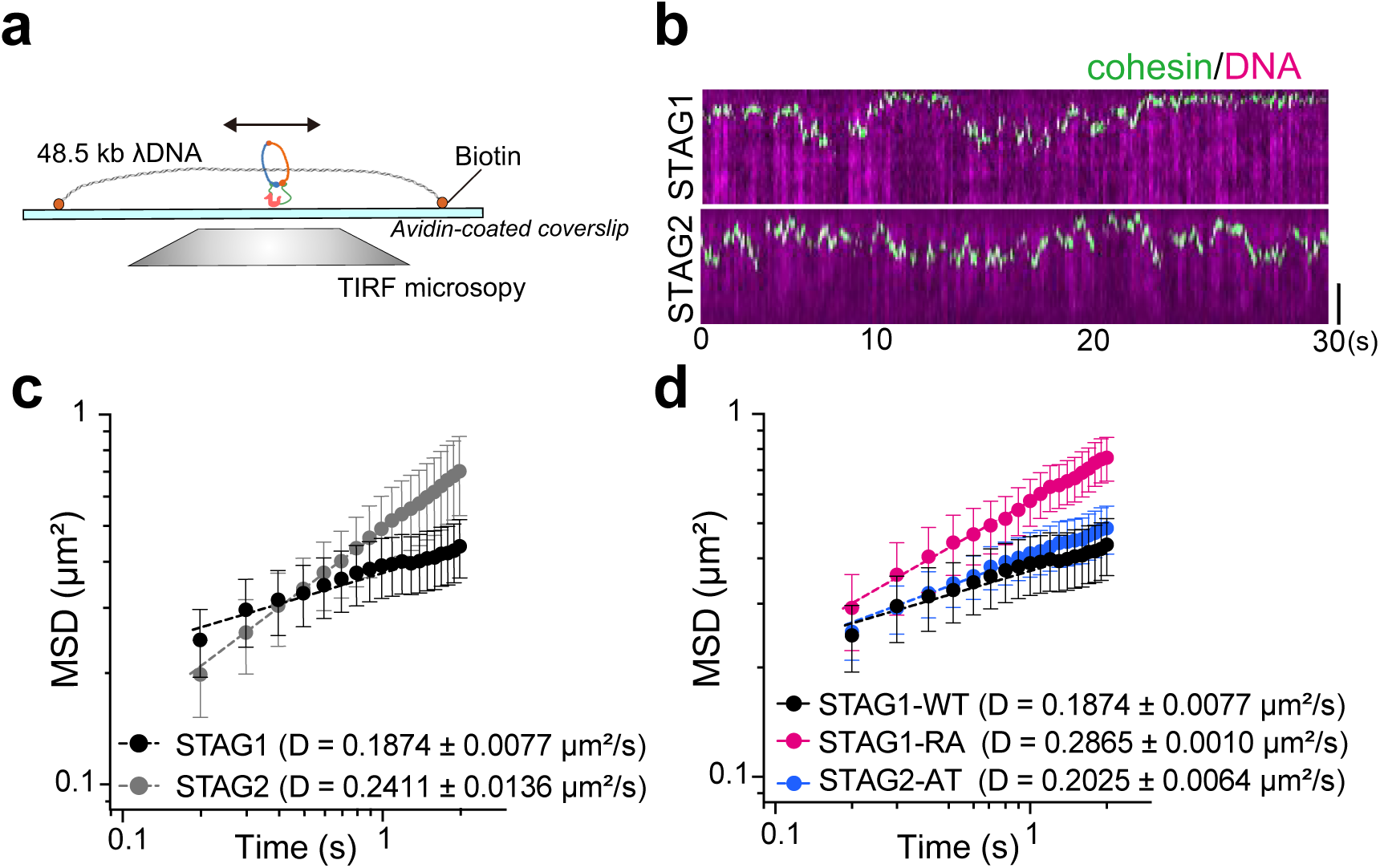
The AT-hook restricts cohesin dynamics on DNA. **a,** Schematic representation of the cohesin translocation assay. Biotin-labeled λ-DNA molecules were tethered in a linear conformation to an avidin-coated coverslip. Cohesin was loaded onto DNA in the presence of NIPBL-Mau2 and ATP, followed by high-salt wash to remove unbound or loosely associated proteins. DNA-bound cohesin particles were tracked at 0.1-second intervals for 30 seconds. **b,** Kymograph of Alexa488 labeled cohesin-STAG1 (upper low) and cohesin-STAG2 (lower low). DNA was labeled with SYTOX Orange. Scale bar: 2 μm. **c,** Mean Square Displacement (MSD) versus time for DNA-bound cohesin-STAG1 and cohesin-STAG2, from which the diffusion (D) coefficient is calculated (MSD ∝ t^α^, α < 1; α = 0.2325 and 0.5291 for cohesin-STAG1 and -STAG2, respectively). The D values were calculated from more than 25 cohesin particles per condition (mean ± SEM). **d,** MSD versus time for DNA-bound cohesin-STAG1-WT, cohesin-STAG1-RA and cohesin-STAG2-AT (MSD ∝ t^α^, α < 1; α = 0.4018 and 0.2626 for cohesin-STAG1-RA and -STAG2-AT, respectively). Data of STAG1-WT, used for comparison, is same as **c**. The D values were calculated from more than 22 cohesin particles per condition (mean ± SEM).

We next asked whether the AT-hook is required for the stability of cohesin-STAG1. We found that the diffusion coefficient was significantly increased in STAG1^RA^ (D = 0.2865 ± 0.0010 μm^2^/s) compared to STAG1^WT^ (D = 0.1874 ± 0.0077 μm^2^/s), whereas STAG2^AT^ exhibited enhanced stability closely resembling STAG1^WT^ (D = 0.2025 ± 0.0064 μm^2^/s, Fig. 4d). These results suggest that AT-hook reinforces DNA-cohesin association after loading on DNA, thereby promoting efficient initiation of loop extrusion.

### The AT-hook stabilizes cohesin on chromatin in G1 but not in G2 phase

We next investigated the physiological function of the AT-hook in living cells. For this purpose, we established HCT116 cell lines in which endogenous STAG1 and STAG2 were tagged with FKBP12^F36V^ and mini-AID degrons, respectively (Fig 5a and Extended Data Fig. 4a,b). Thus, their levels could be reduced below 3% and 5% of those in untreated cells by addition of dTAGV-1 and 5-phenyl-indole-3-acetic acid (5-ph-IAA) respectively^44–46^ (Extended Data Fig. 4c-e), and replaced with exogenously expressed Halo-tagged wild-type STAG1 (STAG1^WT^-Halo) or AT-hook mutated STAG1 (STAG1^RA^-Halo, Fig.5a, Extended Data Fig. 5a-c). We first examined whether the AT-hook is required for the chromatin binding of cohesin. After the degradation of endogenous STAG1 and STAG2, cohesin (Smc1) was not detectable in the chromatin-bound fraction (Extended Data Fig. 5d-f). However, upon induction of either STAG1^WT^ or STAG1^RA^ expression, chromatin-bound cohesin was restored to a level comparable to that of untreated cells in both G1 and G2 phases (Extended Data Fig. 5d-f), indicating that AT-hook is dispensable for the overall chromatin binding of cohesin.

**Fig. 5.**
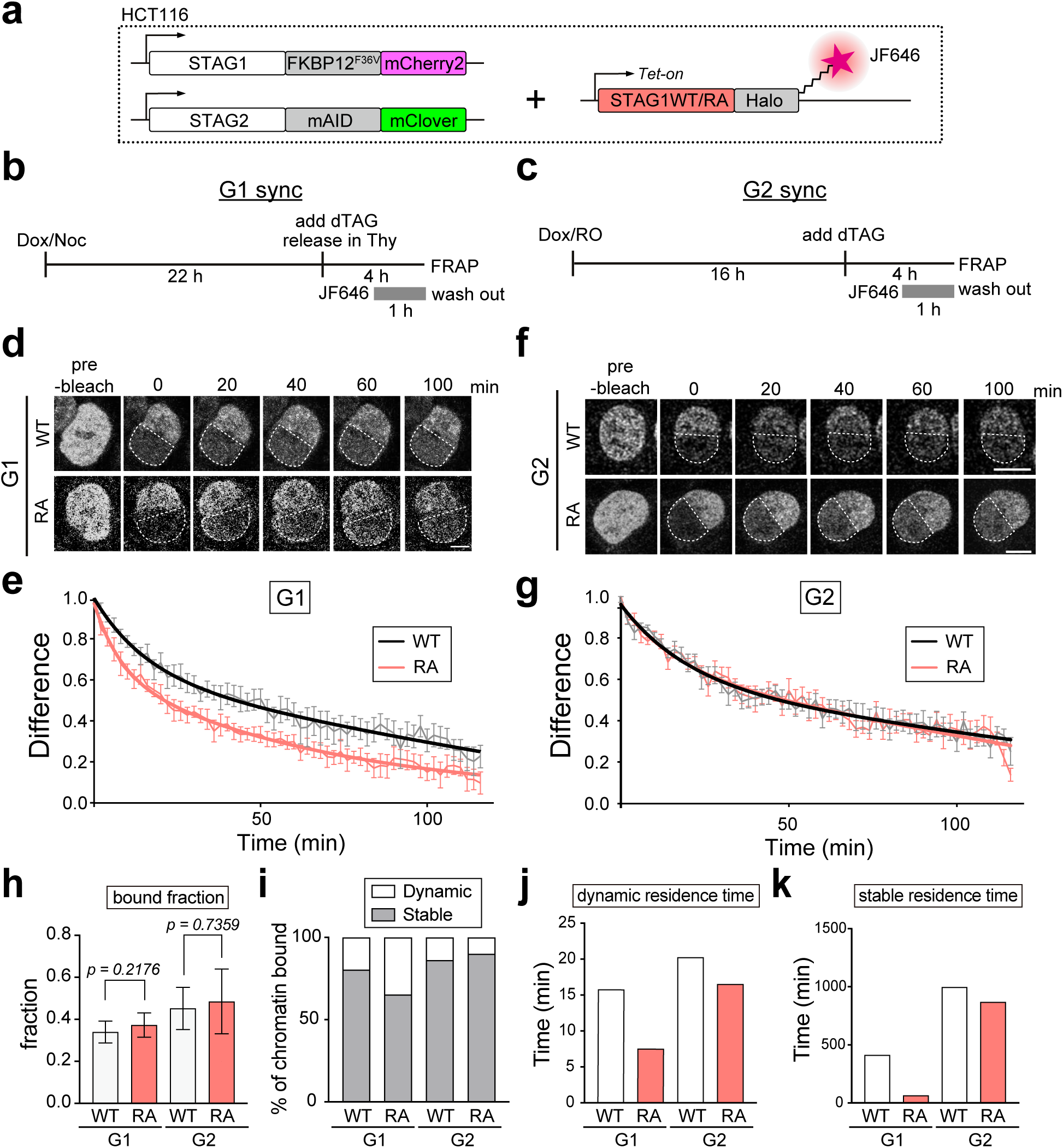
The AT-hook stabilizes cohesin on chromatin in G1 phase. **a,** Schematic representation of the exogenous STAG1^WT^- or STAG1^RA^-Halo-expressing cell lines. The parental cell line was a STAG1/2 double degron cell line. The exogenous constructs were Halo-tagged and labeled with JF646. **b**, **c,** Experimental outline for cell synchronization in G1 or G2 phase for iFRAP analysis. STAG1^WT^- or STAG1^RA^-Halo was labeled with JF646 for 1h and unbound ligand was washed out immediately before the iFRAP experiment. **d,** Representative still images from iFRAP experiments of STAG1^WT^- or STAG1^RA^-Halo-expressing cells in G1. Scale bar, 10 μm. **e,** Normalized fluorescence recovery curves of STAG1^WT^-Halo (black) and STAG1^RA^-Halo (pink) in G1. Solid lines represent bi-exponential curve fits. n = 10 cells per condition (mean ± SEM). **f,** Representative still images from iFRAP experiments of STAG1^WT^- or STAG1^RA^-Halo-expressing cells in G2. Scale bar, 10 μm. **g,** Normalized fluorescence recovery curves of STAG1^WT^-Halo (black) and STAG1^RA^-Halo (pink) in G2. Solid lines represent bi-exponential curve fits. n = 10 cells per condition (mean ± SEM). **h,** Quantification of the chromatin bound fraction of STAG1^WT^-Halo and STAG1^RA^-Halo in G1 and G2. n = 10 cells per condition (mean ± SD). The P values were assessed by a two-tailed Mann-Whitney U test. **i,** Fraction of dynamically and stably bound STAG1^WT^-Halo and STAG1^RA^-Halo determined by bi-exponential fitting of the curves shown in **e** and **g**. **j, k,** Chromatin residence times of the dynamic (**j**) and stable (**k**) fractions of STAG1^WT^-Halo and STAG1^RA^-Halo in G1 and G2 determined by bi-exponential fitting of the curves shown in **e** and **g**.

To analyze cohesin dynamics on chromatin in more detail, we next performed inverse fluorescence recovery after photobleaching (iFRAP). Briefly, after synchronization in G1 and G2 phase, half of the nucleus was photobleached and fluorescence recovery of endogenous STAG1 (labeled with mCherry2) or STAG2 (labeled with mClover) was monitored (Extended Data Fig. 6a,b). The resulting fluorescence curves were fitted using a two-phase exponential model, which allowed us to resolve both a rapidly exchanging *dynamic fraction* and a more stably bound *stable fraction* of STAG1 and STAG2 (Extended Data Fig. 6c,d). Consistent with previous work in human and mouse cells^27,28,47^, cohesin was more stably bound in G2 than in G1, and STAG1 consistently showed greater chromatin stability than STAG2, with slower recovery, higher chromatin-bound fraction, and increased proportion and residence time of the stable population (Extended Data Fig. 6c-h).

We then evaluated the dynamics of exogenously expressed STAG1^WT^- and STAG1^RA^-Halo. Following expression of STAG1-Halo and degradation of endogenous STAG1, Halo-tagged proteins were labeled with JF646 (Fig. 5b,c). In G1 phase, STAG1^WT^ exhibited slow fluorescence recovery, whereas STAG1^RA^ recovered rapidly, indicating reduced stability on chromatin (Fig. 5d,e). Though the overall chromatin-bound fraction was similar between STAG1^WT^ and STAG1^RA^ (Fig. 5h), consistent with western blot analyses (Extended Data Fig. 5f), STAG1^RA^ showed a pronounced reduction in the proportion of the stable fraction and a significant decrease in the residence times of both the dynamic and stable fractions (Fig. 5i-k). Thus, AT-hook is required for stable chromatin association of cohesin during G1 phase.

In contrast to G1 phase, both STAG1^WT^ and STAG1^RA^ were similarly stable in G2 phase (Fig. 5f, g). STAG1^RA^ showed comparable recovery kinetics to STAG1^WT^ (Fig. 5g), and no significant differences were observed between STAG1^WT^ and STAG1^RA^ in terms of bound fraction, stable fraction or residence time during G2 phase (Fig. 5h-k). This finding suggests that AT-hook is not required for stable chromatin association during G2 phase. Given that cohesin is stabilized by Sororin during S/G2 phase to establish sister chromatid cohesion^48^, we assumed that cohesion-establishment mechanism may compensate for the absence of AT-hook. To test this possibility, we performed iFRAP after Sororin depletion. Sororin depletion resulted in severe cohesion defects both in STAG1^WT^ and STAG1^RA^ cells (Extended Data Fig. 7a-c), indicating efficient Sororin knockdown. Under this condition, all STAG1, STAG2, and STAG1^RA^ became more dynamic in G2 phase (Extended Data Fig. 7d-h and 8). These findings suggest that AT-hook-dependent mechanism of cohesin stabilization can be compensated by Sororin-dependent cohesion establishment.

### The AT-hook is not required for sister chromatid cohesion

Because sister chromatid cohesion depends on Sororin even in STAG1^RA^ cells (Extended Data Fig. 7c), we speculated that the AT-hook is dispensable for the sister chromatid cohesion. To test this notion, we compared topological DNA binding activity *in vitro* between cohesin-STAG1^WT^ and -STAG1^RA^, and we found that both types of cohesin exhibited a similar level of this activity (Fig. 6a). We next tested whether the AT-hook is required for sister chromatid cohesion *in vivo* using the double-degron HCT116 cell line (Fig. 5a). Simultaneous degradation of both STAG1 and STAG2 led to dramatic cohesion defects with completely separated sister chromatids (Fig. 6b-d). Remarkably, exogenous expression of either STAG1^WT^ or STAG1^RA^ restored cohesion to a similar extent (Fig. 6d), indicating that AT-hook is dispensable for sister chromatid cohesion. These results suggest that while AT-hook-dependent stabilization of cohesin is required for DNA loop extrusion, it is distinct from the mechanism of sister chromatid cohesion.

**Fig. 6.**
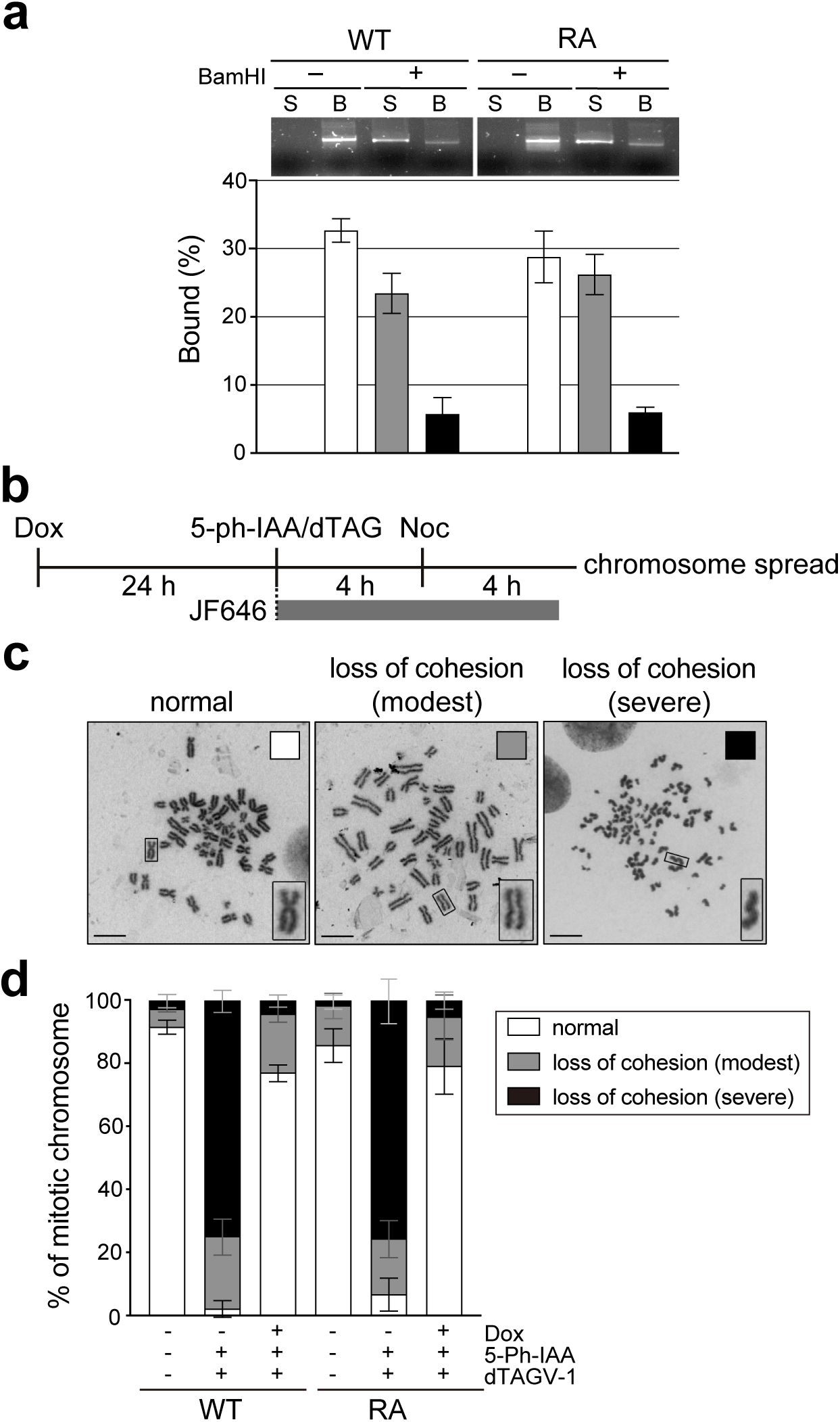
The AT-hook is dispensable for sister chromatid cohesion. **a,** *In vitro* topological DNA binding assay. Restriction enzyme (BamHI) treatment was performed after the cohesin-bound DNA pulldown. Data were collected from 3 independent trials (mean ± SD). Supernatant (S) and beads-bound (B) fractions were shown. **b,** Experimental outline of chromosome spread of STAG1^WT^- or STAG1^RA^-Halo-expressing cells, coupled with degradation of endogenous STAG1/2. **d,** Mitotic chromosomes from STAG1^WT^- or STAG1^RA^-Halo-expressing cells, analyzed by spreading and Giemsa staining. Scale bars, 10 μm **e,** Quantification of cohesion phenotypes shown in **d**. Data were collected from 3 independent trials (mean ± SD).

### STAG1 organizes higher-order chromatin structure through AT-hook motif

We next asked whether AT-hook-dependent loop extrusion mechanism had any impact on higher-order chromatin structure in HCT116 cells. To answer the question, we performed Micro-C in STAG1^WT^- or STAG1^RA^-Halo-expressing cells, following cell cycle synchronization in G1 phase and degradation of endogenous STAG1 and STAG2 (Extended Data Fig. 9a). Consistent with previous observations^49^, chromatin contacts disappeared in the absence of both STAG1 and STAG2 (Fig. 7a (i), (iv) vs (ii), (v)). Exogenous expression of either STAG1^WT^ or STAG1^RA^ restored chromatin interactions but with important differences between the two conditions (Fig. 7a (iii) vs (vi)), including the displacement of the contact density profile at cohesin-mediated loop sizes (0.1-0.5 Mb, shaded in Fig. 7b), indicative of loop enlargement, the reduction in the number of small loops (Extended Data Fig. 9b), weaker TADs, particularly the smaller ones (Fig. 7c and Extended Data Fig. 9c), and the lower insulation score (Fig. 7d) in the STAG1^RA^-Halo-expressing cells. In addition, compartment strength, which rises in cells without endogenous STAG proteins due to the lack of loop-extruding cohesin, is restored upon expression of STAG1^WT^ but not STAG1^RA^ (Fig. 7e). These results suggest that AT-hook not only promotes STAG1-dependent loop extrusion, but also strengthens TAD boundaries.

**Fig. 7.**
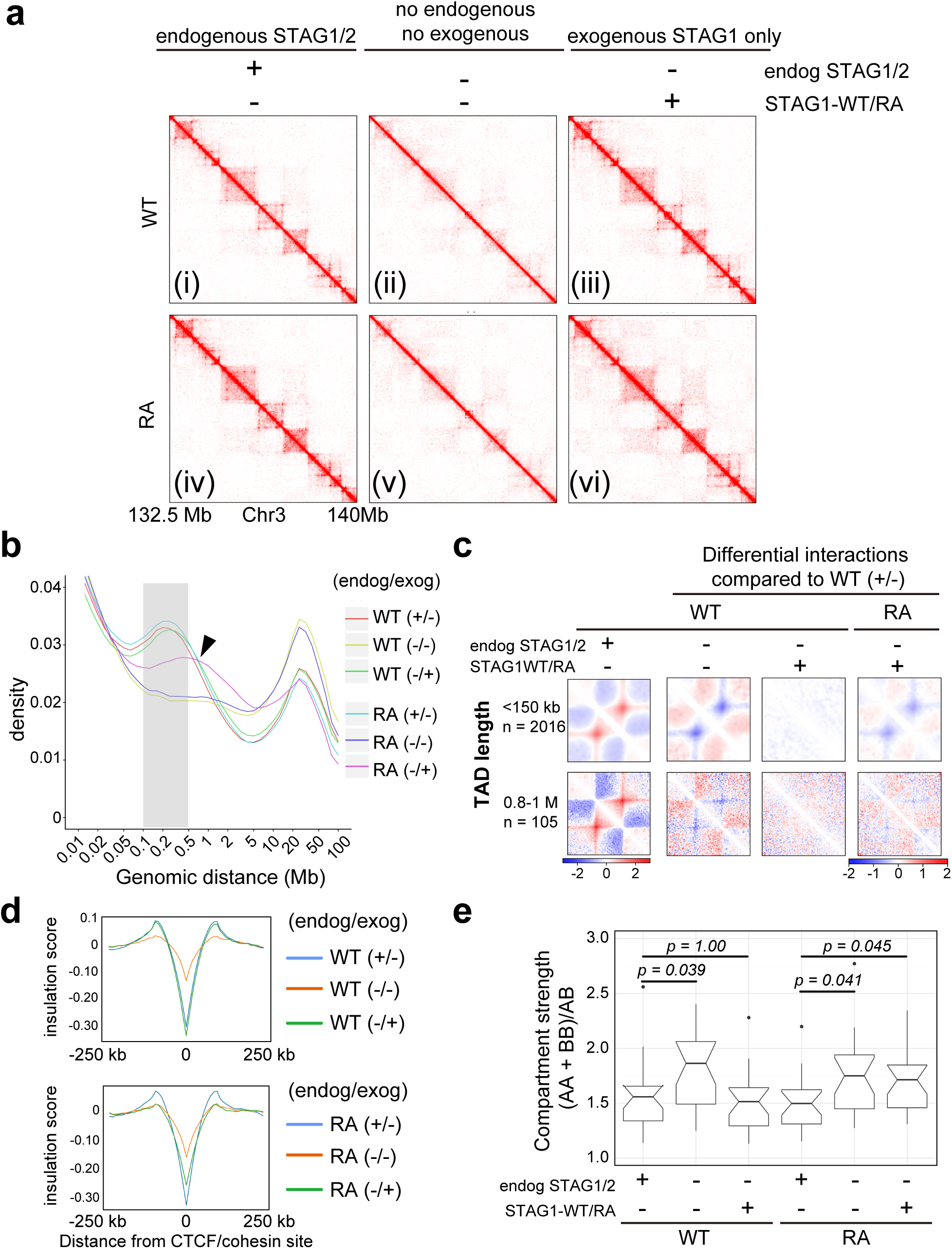
STAG1 organizes higher-order chromatin structure through AT-hook motif. **a**, Micro-C contact matrices for region 132.5-140 Mb in chromosome 3 for the different conditions at 25-kb resolution. The arrowhead and the box highlight differences between cells expressing STAG1^WT^ and STAG1^RA^. **b,** Contact density according to genomic distance. The size range of cohesin-mediated interactions is shadowed. The arrowhead indicates the displacement of the curve towards higher sizes for the STAG1^RA^ cells. **c**, Differential interactions found in the indicated conditions compared to WT in the presence of endogenous STAG1/2 and the absence of exogenous STAG1 (+/-), at TADs of the indicated size described by Rao 2017^49^. STAG1^RA^ expressing cells show severe decreased recovery of TADs of short length (<150 kb) compared to long length (0.8-1 Mb). See **Extended Data Fig.9c** for all TAD sizes. **d**, Insulation score at 10,000 CTCF/Cohesin sites (ChIP-seq data from Porter 2023^67^). **e**, Compartment strength in the different conditions. Boxes represent interquartile range (IQR); the midline represents the median; whiskers are 1.5 x IQR; and individual points are outliers. P values were calculated with Mann-Whitney U test corrected by FDR.

### The AT-hook promotes cohesin-CTCF interaction and SMC3 acetylation

Stable chromatin binding of cohesin depends on Smc3 acetylation at K105/106, catalyzed by Esco1, and on CTCF interaction, providing one of the underlying mechanisms for interphase genome organization^28^. Previous studies have suggested that Smc3 acetylation is promoted by cohesin-STAG1 interaction with CTCF, which may stabilize chromatin-bound cohesin by competing with Wapl^12,27–29,50^. As STAG1^RA^ was more dynamic on G1 chromatin (Fig. 5) and exhibited impaired insulation (Fig. 7d), it is plausible that AT-hook stabilizes cohesin by facilitating Smc3 acetylation and/or interacting with CTCF, thereby strengthening TAD boundaries in G1 phase. To test this possibility, we quantified the level of Smc3 acetylation, in STAG1^WT^- or STAG1^RA^-expressing cells. In these cells, following degradation of endogenous STAG1/2, total Smc3 levels were comparable between STAG1^WT^- and STAG1^RA^-expressing cells whereas acetylated Smc3 was decreased by ∼53% in STAG1^RA^ cells specifically during G1, but not G2 phase (Fig. 8a-c). Alongside this reduction in acetylated Smc3, the amount of CTCF co-immunoprecipitated with cohesin was decreased by approximately 40% in STAG1^RA^-expressing cells (Fig. 8d-f). Together, these results demonstrate that the AT-hook is required for stable chromatin association of cohesin-STAG1 by both reinforcing DNA binding and promoting Smc3 acetylation and cohesin-CTCF interaction.

**Fig. 8.**
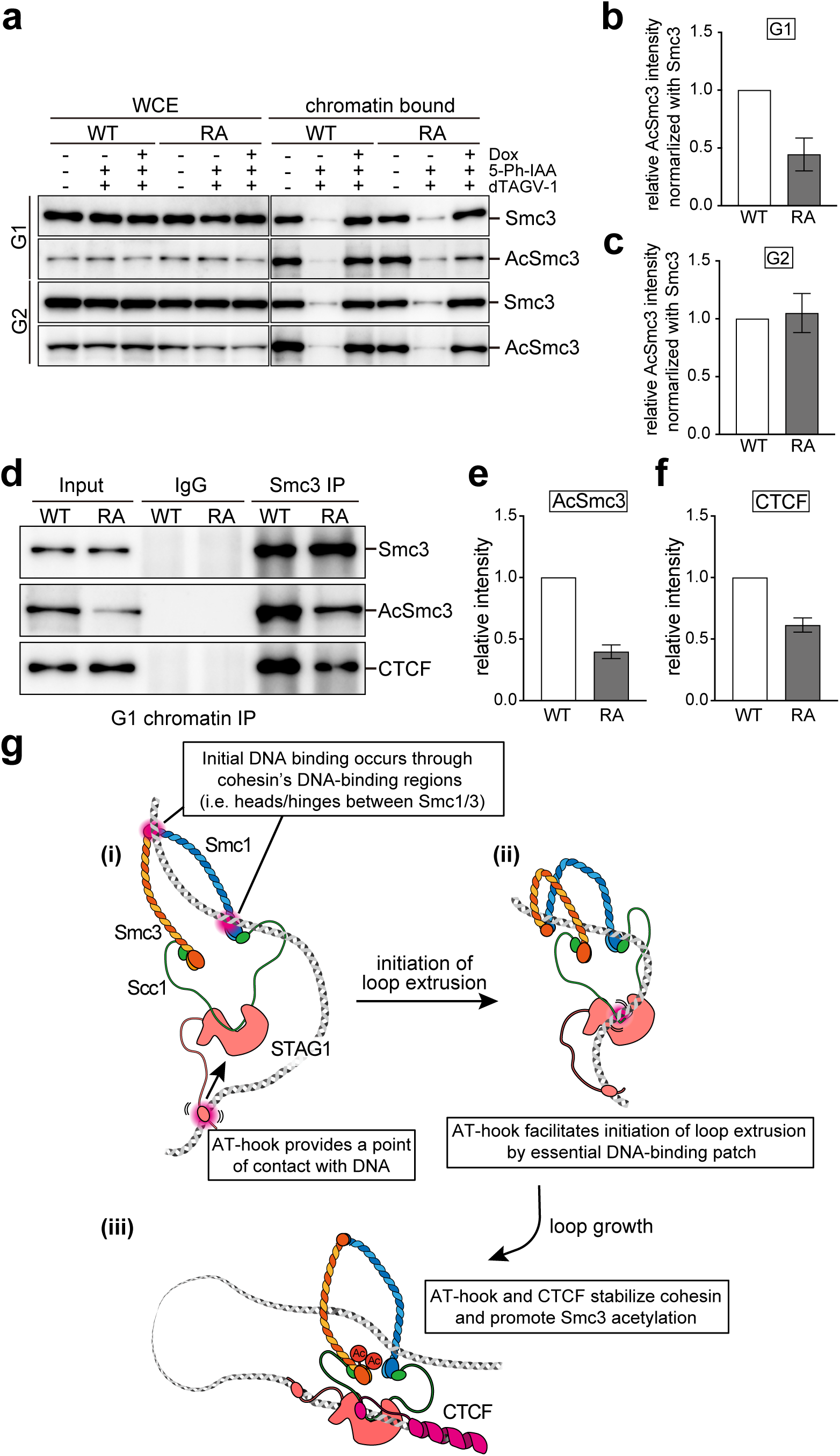
The AT-hook mutation diminishes Smc3 acetylation and interaction of cohesin with CTCF on chromatin. **a,** Western blot analysis of Smc3 and acetylated Smc3 levels of whole-cell extracts (WCE) and chromatin-bound fractions from STAG1^WT^- or STAG1^RA^-Halo-expressing cells synchronized in G1 and treated with the indicated components. **b,c,** Quantification of the AcSmc3 signal of STAG1^WT^- or STAG1^RA^-Halo-expressing cells in G1 (**b**) and G2 (**c**) relative to Smc3 in **a**. The AcSmc3/Smc3 ratio was normalized to that of STAG1^WT^cells. Data were collected from 3 independent trials (mean ± SD). **d,** Immunoprecipitation of chromatin fractions from STAG1^WT^- or STAG1^RA^-Halo-expressing cells synchronized in G1, following degradation of endogenous STAG1 and STAG2. **e, f,** Quantification of immunoprecipitated AcSmc3 (**e**) and CTCF (**f**) signals relative to Smc3 in **d**. The AcSmc3/Smc3 and CTCF/Smc3 ratios were normalized to that of STAG1^WT^. Data were collected from 3 independent trials (mean ± SD). **g**, Schematic model illustrating a possible mechanism by which the AT-hook motif facilitates initiation of DNA loop extrusion and cohesin stabilization in cell. (i) Cohesin initially binds DNA through DNA binding region within the complex, and the AT-hook provides a flexible contact point, thereby stabilizing cohesin. (ii) AT-hook stabilizes cohesin and promotes initiation of DNA loop extrusion, which is driven by the essential DNA-binding patch within the HEAT repeat domain of STAG1. (iii) Subsequent loop growth and maintenance occur independently of the AT-hook. In the cellular context, the DNA loop extrusion activity of cohesin-STAG1, which in turn increases the probability of cohesin engaging with CTCF and being anchored at CTCF-binding sites. This anchoring promotes Smc3 acetylation, further stabilizing loop-extruding cohesin and reinforcing cohesin-dependent 3D genome architecture.

## Discussion

In this study, we identify an AT-hook motif in the N-terminus of STAG1 as a major structural determinant of the different contributions of cohesin-STAG1 and cohesin-STAG2 to 3D genome organization.

### The AT-hook motif distinguishes cohesin-STAG1 and -STAG2 as loop extruders but not as cohesive complexes

Our *in vitro* analyses revealed intrinsic differences in loop extrusion between cohesin-STAG1 and cohesin-STAG2. Despite comparable ATPase activity and topological DNA entrapment, cohesin-STAG1 exhibited markedly higher loop extrusion activity and stability in both DNA and chromatin context. This enhancement depends critically on the STAG1-specific AT-hook motif, absent in STAG2. Mutation of the AT-hook diminished loop extrusion and increased dynamics on DNA/chromatin, whereas insertion into STAG2 promoted loop extrusion and cohesin stabilization, indicating that AT-hook is a key structural element for efficient loop extrusion and stable DNA association by STAG1. Although AT-hook can bind the DNA minor groove and increase the affinity of monomeric STAG1^36,37^, our single-molecule assays showed no effect on initial binding efficiency of cohesin-STAG1, suggesting compensation by other DNA-binding elements in cohesin such as Smc1/3 or cohesin loader NIPBL^38–40^ . Indeed, biochemical study suggested that the initial DNA binding occurs at the hinge or head domains of Smc1/3^39^. Thus, the AT-hook specifically stabilizes cohesin once bound to DNA, thereby facilitating the initiation of loop extrusion (Fig. 8g (i),(ii)), but it is dispensable for topological entrapment -highlighting mechanistic distinctions between loop extrusion and cohesion.

Cohesin is topologically associated with replicated DNA to maintain sister chromatid cohesion^51^. As the AT-hook is dispensable for topological binding to DNA (Fig. 6a) and sister chromatid cohesion (Fig. 6d), these “cohesive cohesin” molecules are likely present on chromatin independently of the AT-hook and stabilized by Smc3 acetylation and Sororin (Fig. 8 and Extended Data Fig. 8). Because STAG1^RA^ is still more stable on chromatin in G2 phase than STAG2, differences in chromatin-bound stability between STAG1 and STAG2 cannot be solely attributed to AT-hook, suggesting that additional factors contribute to either stabilize STAG1 or destabilize STAG2. Although previous study has suggested that the AT-hook is required for telomere cohesion^36^, this could be due to either: 1) overexpression of STAG1 simply leading to over cohesion, under which condition STAG1^RA^ instability becomes more evident as a cohesion phenotype; or 2) telomere cohesion is regulated by distinct mechanisms from those regulating other genomic regions.

### DNA loop extrusion-dependent stabilization of cohesin and chromatin structure

Our results in human cells demonstrated that the AT-hook motif is required for the stable association of STAG1 with chromatin in G1 phase. Cohesin complexes carrying STAG1^RA^ are less dynamic than cohesin-STAG2, implying that additional elements contribute to the different behavior of the two complexes also in G1. Our Micro-C analyses showed that AT-hook mutation impairs TADs and chromatin loops in human cells (Fig. 7). It also diminished Smc3 acetylation and the association of cohesin with CTCF on chromatin (Fig. 8). Since both Smc3 acetylation and CTCF promote stabilization of cohesin-STAG1^28^, one possible mechanism of AT-hook-dependent cohesin stabilization on chromatin is that AT-hook facilitates loop formation, promotes cohesin association with CTCF, and thereby stabilizes TADs. Single-molecule analysis showed that CTCF can halt loop extruding cohesin at its binding site^52^. Several studies support the notion that the stalling of loop extrusion by CTCF constitutes a fundamental process in the formation of 3D genome structure^53,54^. Even though the YDF motif in the N-terminal region of CTCF interacts with a region shared by STAG1 and STAG2^12^, and both cohesin colocalize with CTCF, cohesin-STAG1 is more persistent at CTCF sites after NIPBL or CTCF knock down^55^ and preserves better TAD boundaries^27,28^. Given our findings that STAG1 exhibits higher loop extrusion activity and that the loop extrusion-deficient STAG1^RA^ shows diminished CTCF association, we propose that AT-hook-dependent loop extrusion stabilizes cohesin to traverse longer distances, thereby increasing the probability to reach and be stabilized at CTCF-bound loci, whereas impaired loop extrusion reduces such encounters and diminishes CTCF anchoring. Previous studies have shown that CTCF depletion decreases Smc3 acetylation^28^, suggesting that CTCF not only anchors cohesin but also promotes Smc3 acetylation. Thus, loop extrusion may facilitate CTCF association, which in turn enhances Smc3 acetylation, together reinforcing cohesin stabilization (Fig. 8g (iii)). However, because Smc3 acetylation was more severely decreased than CTCF association in STAG1^RA^, additional mechanisms might be involved. One possibility is that the instability of STAG1^RA^ on chromatin restricts its chance to interact with Esco1 or favors its interaction with the cohesin deacetylase HDAC8.

### Is DNA loop extrusion a conserved mechanism for chromatin organization?

Recent models emphasize HEAT-repeat subunits play crucial roles in loop extrusion by SMC complexes, anchoring or reeling DNA through its DNA-binding ability^14,17,18,56^. Because a DNA-binding patch essential for loop extrusion is present in the middle region of STAG1/2 and its yeast ortholog Scc3^14,15^, it could be one of the ancestral loop extrusion machineries. Notably, STAG1^RA^ still retained minimal LE activity, suggesting that AT-hook may facilitate loop extrusion mediated by DNA-binding patch in the middle region. AT-hook is located in the N-terminal IDR, which likely allows for conformational flexibility. Because loop extrusion requires dynamic conformational changes in cohesin^14,18^, the AT-hook may stabilize cohesin-DNA interactions independently of cohesin conformation (Fig. 8g (i)), thereby promoting initiation of loop extrusion by essential DNA-binding patch (Fig. 8g (ii)). Accordingly, cohesin-STAG2, which lacks the AT-hook, may initiate extrusion less frequently, though subsequent loop growth and loop stabilization appear comparable between STAG1 and STAG2, because both possess conserved DNA binding patch. Together, these findings identify that AT-hook as a STAG1-specific loop extrusion enhancer, facilitating initiation by stabilizing cohesin-DNA interactions in cooperation with essential DNA-binding patch in both paralogs.

Although the AT-hook is unique to vertebrate STAG1, organisms with a single STAG (Scc3), such as *Drosophila*, *C. elegans*, and yeasts, harbor similar lysine/arginine-rich clusters in the N-terminal region (Extended Data Fig. 10). If these KR-clusters act analogously to the AT-hook, ancestral cohesin complexes may exhibit activities resembling cohesin-STAG1. Indeed, yeast cohesin has shown to extrude DNA loop *in vitro*^15,18^. Consistent with this idea, TAD-like domains and loop-like structures are observed in invertebrates too (e.g. *S. cerevisiae*^15,57^, *S. pombe*^58^, *Drosophila*^59^, and *C. elegans*^60^). However, recent study showed that meiotic cohesin component STAG3, which also has KR-clusters in its N-terminal IDR (Extended Data Fig. 10), exhibits defect in insulation and CTCF co-localization in mouse germline stem cells^61^, implying that STAG3 could be a weak loop extruder. Another study in yeast demonstrated that intact chromatin structure can be formed even by loop extrusion-defective mutant^15^. Given these observations, the conserved mechanisms underlying genome 3D folding across diverse cell types and organisms cannot be explained solely by the loop extrusion associated with KR clusters. Rather, loop extrusion may cooperate with other chromosomal factors to form higher-order chromatin structures, and the degree of contribution could vary depending on the organism and cell type. If so, the emergence of STAG2, lacking the AT-hook, in other words “weak loop extruder”, may allow more fine-tuned and context-dependent gene regulation, including tissue-specific gene activation and super-enhancer control^29–31^. Mutation in STAG2-cohesin may disrupt these fine-tuned gene regulations and cause genetic disorders including cancers^62^.

Taken together, the AT-hook promotes stable chromatin association of cohesin-STAG1 by acting as a DNA-binding module and as a loop extrusion enhancer. It stabilizes loop anchors by promoting CTCF interaction and Smc3 acetylation, thereby supporting higher-order chromatin structures such as TADs. Because AT-hook-like KR-clusters are conserved in yeast Scc3, the N-terminal DNA-binding region of STAG1 not only distinguishes it from STAG2 but also represents a fundamental molecular feature of cohesin function in genome organization across evolutionary divergence.

## Methods

### Cell lines

HCT116 cells (CCL-247) served as the parental cell line for all genome editing experiments. All cell lines were cultured in McCoy’s 5A medium (Thermo Fisher) supplemented with 10% fetal bovine serum (Gibco), 100 U/ml penicillin, and 100 μg/ml streptomycin (Wako) at 37 °C in a humidified incubator with 5% CO₂. To generate STAG1-FKBP12^F36V^-mCherry2/STAG2-mAID-mClover knock-in cell lines, cells were transfected in six-well plates with CRISPR/Cas9 and donor plasmids using Lipofectamine 3000 (Thermo Fisher). The FKBP12^F36V^-mCherry2 and mAID-mClover tags were inserted at the C-terminus of endogenous STAG1 and STAG2 respectively, using the following guide DNAs: STAG1-C (forward: 5′-caccggtcatcatttttctagggatt-3′, reverse: 5′-aaacaatccctagaaaaatgatgac-3′), and STAG2-C (forward: 5′-caccgtctctctctcattaggttct-3′, reverse: 5′-aaacagaacctaatgagagagaga-3′). Three days post-transfection, cells were selected with 0.7 mg/ml G418 (Roche) and 0.1 mg/ml HygroGold (InvivoGen). After 10-12 days of selection, resistant colonies were picked and expanded in 48-well plates. To express OsTIR1^F74G^ under the CMV promoter, lentiviral transduction was performed. Viral supernatants were produced in HEK293T cells and applied to HCT116 cells in the presence of 8 μg/ml polybrene, followed by selection with 400 μg/ml Zeocin (InvivoGen). Plasmids encoding STAG1 (provided by K. Yokomori) were used to mutate the AT-hook in STAG1. A doxycycline-inducible Halo-tagged STAG1 transgene encoding either wild-type (STAG1^WT^) or an AT-hook mutant (STAG1^RA^) was introduced by lipofection using Lipofectamine 3000 and selected with 6 μg/ml blasticidin S (Wako). To induce the expression of exogenous STAG1^WT^- or STAG1^RA^-Halo and degradation of endogenous STAG1 and STAG2, cells were treated with 2 μg/ml doxycycline hycrete (sigma) for 24 h, 500 ng/ml dTAGV-1 (Wako) for 4h and 100 nM 5-Ph-IAA (sigma) for at least 4 h.

### Protein purification

Recombinant human trimeric cohesin (Scc1^Halo^, Smc3^FLAG^, Smc1), tetrameric cohesin (10×His-tagged STAG1/STAG2/STAG1^RA^/STAG2^AT^, Scc1^Halo^, Smc3^FLAG^ Smc1) or ^FLAG-Halo^NIPBL^10His^-Mau2 were expressed in *Spodoptera frugiperda* (Sf21) cells using the baculovirus expression system. Coding sequences of each protein was cloned into pAceBac1 vector and introduced into Sf21 cells via baculovirus infection. For cohesin purification, infected cell pellets (∼1 ml) were resuspended in 3 ml of CytoBuster reagent (Millipore) supplemented with cOmplete EDTA-free protease inhibitor cocktail (Roche), 0.2% Tween-20, and 0.5 mg/ml PMSF, followed by incubation at 4 °C for 45 min. After centrifugation at 15,000 rpm for 10 min, the supernatant was incubated with anti-FLAG M2 affinity gel (Sigma-Aldrich) at 4 °C for 2.5 h.

For trimeric cohesin, the resin was washed with Buffer A (25 mM Tris-HCl pH 8.0, 100 mM NaCl, 10% glycerol, 0.5% Tween-20), and bound proteins were labeled with 16 μM HaloTag Alexa Fluor 488 ligand (Promega) for 1 h at 4 °C. Labeled proteins were eluted with Buffer A supplemented with 0.2 mg/ml 3×FLAG peptide and dialyzed into Buffer A.

For tetrameric cohesin, FLAG-eluted proteins were incubated with Ni²⁺-NTA agarose resin (Thermo Fisher) at 4 °C for overnight. Bound cohesin was incubated with 16 μM HaloTag Alexa Fluor 488 ligand for 1 h at 4 °C. Labeled complexes were eluted with Ni Buffer B (25 mM potassium phosphate pH 7.4, 150 mM KCl, 200 mM imidazole, 10% glycerol), and peak fractions were dialyzed into Buffer A.

For NIPBL-Mau2, Sf21 cells expressing ^FLAG–Halo^NIPBL^10His^-Mau2 (∼1 ml pellet) were resuspended in 3 ml of purification buffer 1 (25 mM sodium phosphate, pH 7.5, 500 mM NaCl, 5% glycerol, 0.05% Tween-20) supplemented with 1 mM PMSF, 2 mM benzamidine, 10 μl/ml aprotinin, and cOmplete EDTA-free protease inhibitors. Cells were lysed by dounce homogenization (20 strokes) and cleared by centrifugation at 15,000 rpm for 45 min. The supernatant was incubated with 150 μl anti-FLAG resin at 4 °C for 2.5 h with rotation. Resin was washed three times with purification buffer 2 (25 mM sodium phosphate, pH 7.5, 150 mM NaCl, 5% glycerol, 0.01% Tween-20), and bound proteins were eluted with 0.1 mg/ml 3×FLAG peptide for 2 h at 4 °C. Eluates were loaded onto a HisTrap HP column (GE Healthcare), washed with buffer containing 22.5 mM imidazole, and eluted with 300 mM imidazole. Peak fractions (∼2 ml) were pooled and concentrated to ∼200 μl using Vivaspin 20 ultrafiltration units (100 kDa MWCO; Sartorius).

### *In vitro* topological DNA binding assay

Relaxed circular DNA was prepared by incubating covalently closed circular pBluescript KSII (+) with *E. coli* topoisomerase I (New England BioLabs) at 37 °C for 20 min followed by collum purification. For *in vitro* topological DNA binding assay, 50 nM FLAG-Halo-NIPBL-10×His-Mau2 was mixed with 3.3 nM relaxed circular DNA in CL1 buffer (35 mM Tris-HCl, pH 7.5, 1 mM TCEP, 25 mM NaCl, 1 mM MgCl₂, 15% glycerol, 0.003% Tween-20) and incubated on ice for 5 min. Then, 150 nM cohesin was added and incubated for an additional 5 min. The loading reaction was initiated by adding 0.5 mM ATP and incubating at 37 °C for 3 h. The reaction mixture was supplemented with 125 μl of CP buffer (35 mM Tris/HCl, pH 7.5, 0.5 mM TCEP, 500 mM NaCl, 10 mM EDTA, 5% glycerol, 0.35% Triton X-100) and incubated at 37°C for 5 min followed by incubation on ice for further 5 min. Then, 15 μl slurry of anti-Smc3-antibody-coated, protein-G-conjugated magnetic beads (Dynal Inc.) were added into reaction mixture and rotated at 4°C for o/n. The magnetic beads were washed three times with CW1 buffer (35 mM Tris/HCl, pH 7.5, 0.5 mM TCEP, 750 mM NaCl, 10 mM EDTA, 5% glycerol, 0.35% Triton X-100), and then once with CW2 buffer (35 mM Tris/HCl, pH 7.5, 0.5 mM TCEP, 100 mM NaCl, 0.1% Triton X-100). Beads were resuspended in 15 μl elution buffer (10 mM Tris-HCl, pH 7.5, 1 mM EDTA, 50 mM NaCl, 0.75% SDS, 1 mg/ml proteinase K) and incubated at 50 °C for 1 h. Recovered DNA was analyzed by 1% agarose gel electrophoresis in 1× TAE, stained with SYBR Gold (Invitrogen), and quantified using Fiji software.

To linearize entrapped DNA, cohesin-DNA was retrieved by anti-Smc3 immunoprecipitation as described above, except that EDTA was omitted from the CP and CW buffers. Beads were washed with RE buffer (35 mM Tris-HCl, pH 7.5, 0.5 mM TCEP, 100 mM NaCl, 5 mM MgCl₂, 0.1 mg/ml BSA, 0.1% Triton X-100), and incubated with 20 U BamHI (Takara) in 10 μl RE buffer at 10 °C for 3 h. Salt concentration was adjusted to 500 mM NaCl in a total volume of 15 μl, followed by incubation on ice for 5 min. Supernatant and beads-bound DNA fractions were analyzed as described above.

### Coverslip preparation for single-molecule imaging

Single-molecule imaging was performed in cruciform flow-cell channels. Coverslips (22 × 22 mm, No. 1-S; Matsunami) were cleaned by sonication in 1 M KOH for 20 min, rinsed thoroughly with water, and sonicated again in methanol. Then, coverslips were incubated in 2% (v/v) 3-aminopropyltriethoxysilane (Sigma-Aldrich) for 1-3 nights at room temperature.

For surface passivation, coverslips were treatedwith 100 mM NaHCO₃ containing 100 mg/ml methoxy-PEG-succinimidyl valerate (mPEG-SVA-5K; Laysan Bio) and biotin-PEG-succinimidyl valerate (Bio-PEG-SVA-5K; Laysan Bio) at room temperature for o/n. Coverslips were washed gently with water, air-dried, and stored at -80 °C until use. Before imaging, coverslips were incubated with 80 μM streptavidin (Sigma-Aldrich) in ELB++ buffer (10 mM HEPES-KOH, pH 7.7, 50 mM KCl, 2.5 mM MgCl₂, and 1 mg/ml BSA (Sigma-Aldrich)) for at least 30 min at room temperature. Excess streptavidin was removed by washing with water.

### Flow-cell assembly

Cruciform flow-cell channels (2 mm wide) were assembled by sandwiching double-sided tape (Grace Bio Labs) between glass slides and PEG-coated coverslips. Inlet and outlet tubes (Natsume, KN-392-1-SP-19 and KN-392-1-SP-55, respectively) were attached to each channel. Solution flow was controlled by withdrawing with a syringe pump (Pump 11 Elite; Harvard Apparatus) connected to the outlet tubing. Prior to use, channels were washed with ELB++ buffer.

### Biotinylation of λDNA and DNA tethering

Biotinylated λDNA (New England Biolabs) was prepared by ligating biotinylated oligonucleotides to both cos sites. λDNA and two biotinylated oligonucleotides (oligo #1: 5′-GGGCGGCGACC[biotinylated-T]-3′; oligo #2: 5′-AGGT[biotinylated-T]CGCCGCCC-3′) were first phosphorylated with T4 polynucleotide kinase (New England Biolabs) in the presence of ATP at 37 °C for 3 h. To label one end, oligo #1 was annealed to λDNA in T4 ligation buffer (New England Biolabs) by gradually cooling from 65 °C to 10 °C, followed by ligation with T4 DNA ligase (New England Biolabs) at 16 °C for 2 h. Oligo #2 was then annealed and ligated to the other end using the same procedure.

For DNA loop extrusion assay, biotinylated λDNA was injected into the perpendicular channel of the flow-cell at a final concentration of 11.5 pM in DNA buffer (20 mM Tris-HCl pH7.5, 150 mM NaCl, 0.25 mg/ml BSA) containing 22 nM SYTOX Orange (Thermo Fisher Scientific), and tethered onto coverslips under the flow at 2 μl/min for 5-10 min. Unbound λDNAs were washed out with imaging buffer (50 mM Tris-HCl pH7.5, 50 mM NaCl, 3 mM MgCl_2_, 5 mM ATP, 5% glycerol, 0.25 mg/ml BSA, 22 nM SYTOX Orange) flow at 10 μl/min for 2 min. To observe U-shape DNA, the flow direction was switched from the perpendicular to the horizontal channel at a flow rate of 30 μl/min.

For cohesin translocation assays, biotinylated λDNA was introduced into the horizontal channel of the flow cell using the same conditions as the loop extrusion assay, except for a flow rate of 50 μl/min during DNA tethering.

### TIRFM observation of DNA loop extrusion and cohesin translocation

For DNA loop extrusion assay, cohesin and NIPBL-Mau2 were simultaneously introduced into U-shape DNA-tethered flow cells with imaging buffer at a flow rate of 30 μl/min. Cohesin was introduced at 1.5 nM and NIPBL-Mau2 was introduced at 0.75 nM. SYTOX Orange-stained DNA were visualized using a 100 μW 561-nm laser (Coherent Sapphire LPX 561±2, 480 mW) and filmed in every 5 s for 20 min. All assays were performed at room temperature.

For cohesin translocation assay,1.5 nM of cohesin and 2 nM of NIPBL-Mau2 were simultaneously introduced into linearly-tethered DNA with imaging buffer at a flow rate of 10 μl/min and incubated for 15-20 min. After incubation, the flow cell was washed with high salt buffer (50 mM Tris-HCl pH7.5, 500 mM NaCl, 3 mM MgCl_2_, 5 mM ATP, 5% glycerol, 0.25 mg/ml BSA, 22 nM SYTOX Orange) at a flow rate of 10 μl/min for 2 min. Flow was then halted and cohesin particles on DNA were filmed in every 100 ms for 30 s. Cohesin^A488^ and SYTOX Orange-stained DNA were visualized using a 350 μW 488-nm laser (Coherent Sapphire LP 488±2, 150 mW) and 100 μW 561-nm laser (Coherent Sapphire LPX 561±2, 480 mW) respectively. All assays were performed at room temperature. Based on the experimentally obtained trajectory of 25 particles, we determined the mean square displacement (MSD) and diffusion coefficient (*D*) The MSD was calculated as

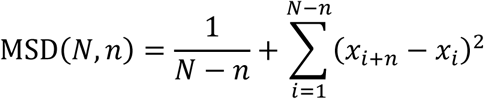

where N is the total number of frames (N = 300) and MSD values were plotted as a function of time lag (*n*Δ*t*) where Δ*t* is the time interval between frames (Δ*t* = 0.1 s). The MSD-time dependence showed subdiffusive behavior, as indicated by a slope less than 1 in log-log plots (Fig.4c,d). Diffusion coefficients (*D*) and anomalous exponents (*α*) were obtained by fitting the MSD data to the anomalous diffusion equation

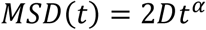

Image acquisition and particle tracking were performed using NIS-Elements (Nikon), MSD analysis and Curve fitting was performed by Prism9 software.

### Bleaching assay

For the bleaching assay to detect single-molecule intensities of cohesin^A488^, 1 nM cohesin^A488^ was non-specifically attached to coverslip. The fluorescently labeled proteins were then bleached by a 3.5 mW 488-nm laser, the images were taken every 0.5 s, and the bleaching step was detected. The step size was determined by calculating the difference in average intensity of five flames before and after the signal was bleached.

### ATPase assay

For ATPase assay, 200 nM of cohesin complexes were incubated in reaction buffer (35 mM Tris/HCl pH 7.5, 1 mM TCEP, 25 mM NaCl, 1 mM MgCl_2_, 0.003 % Tween20, 0.1 mg/ml BSA, 10 nM [γ-32P]-ATP, 25 μM non-radiolabeled ATP) supplemented with 150 nM of NIPBL-Mau2 and 10 nM of λDNA. Reactions were incubated at 37 °C and stopped by adding 0.3 M EDTA. Reaction products were separated on polyethyleneimine plates (Merck) by thin-layer chromatography using 0.75 M KH_2_PO_4_ (pH 3.4) and analyzed by phosphor imaging with a Typhoon FLA9000 Scanner (GE healthcare) and quantified with Fiji software.

### Cell cycle synchronization, FACS and RNAi

For synchronization in G1 phase, cells were treated with 660 nM nocodazole for 22 h and released into media containing 2 mM thymidine for 8 h. For synchronization in G2 phase, cells were treated with 10 μM RO3306 for 20 h.

All experiments using STAG1^WT^- and STAG1^RA^-Halo-expressing cells except iFRAP were conducted coupled with fluorescence-activated cell sorting (FACS) to normalize expression level of exogenous constructs. For FACS, cells were labeled with 100 nM HaloTag ligand Janelia Fluor®646 (JF646; Promega) for 6 h, washed with PBS, harvested by trypsinization, and resuspended in PBS. FACS was performed on SONY SH800HS using 640 nm laser for detection STAG1^WT^- or STAG1^RA^-Halo^JF646^. JF646 positive cells were gated to exclude dead cells and doublets and sorted into McCoy’s 5A medium supplemented with 10 % FBS.

For cell cycle analysis, synchronized cells were harvested by trypsinization and washed with PBS. Cells were resuspended with 1 ml of PBS and fixed by adding 5 ml of 100 % EtOH. After overnight incubation at 4 °C, cells were resuspended with 1 ml PBS containing 1 μg/ml DAPI (Dojindo) and 0.1 % TritonX-100. DNA content was analyzed using 405 nm laser. Data were analyzed using SONY cell sorter software and Floreada.io.

For RNAi experiments, siRNAs were pre-mixed with RNAiMax (Thermo Fisher) according to the manufacturer’s instructions and incubated in the media for 24 h. Sense sequences of siRNA oligos; GCCUAGGUGUCCUUGAGCUtt and Control (GL2) siRNA ; CGUACGCGGAAUACUUCGAtt.

### Live cell imaging and iFRAP

For live cell imaging, cells were cultured in 8-well glass bottom chamber (IWAKI) and labeled with 100 nM HaloTag ligand JF646 for 6 h and 5 μg/ml Hoechst33342 for 10 min prior to imaging. Images were taken on Nikon Ti2-E microscope equipped with a confocal unit CSU-W1 (Yokogawa), an EMCCD camera iXon Ultra 888 (Andor), and 40x Plan-Fluor (NA 1.30) objective (Nikon).

Inverse fluorescence recovery in photobleaching (iFRAP) experiments were performed on cells synchronized in G1 or G2 phase as described above. Cells were imaged on LX83-FV3000 (OLYMPUS) confocal microscope using a 60x Plan-Apochromat objective (PLAPON60XOSC2). For iFRAP of STAG1-FKBP-mCherry2 and STAG2-mAID-mClover, selected 10 cells were bleached by 561 nm laser for STAG1 and 488 nm for STAG2 and leaving a half area of nuclei unbleached. After the bleach, images were taken every 2 min for 120 min. For iFRAP of STAG1^WT^ or STAG1^RA^, cells were treated with 2 ug/ml doxycycline hycrete (sigma-aldrich) for 24 h and 500 ng/ml dTAGV-1 for 4h prior to imaging. After JF646 labeling of STAG1^WT^- or STAG1^RA^-Halo for 1 h at 100 nM, cells were washed with PBS and medium was changed to prewarmed McCoy’s 5A medium without JF646, and Hoechst33342 was added at a final concentration of 5 μg/ml to visualize DNA, 30 min before imaging. Selected 10 cells were bleached by 640 nm laser and leaving a half area of nuclei unbleached. Fluorescence intensities in bleached and unbleached regions were quantified using Fiji and normalized. Cell movements were corrected using the Template Matching plugin in Fiji. The decay of fluorescence intensity over time was fitted to a two-phase exponential function using Prism 9 (GraphPad):

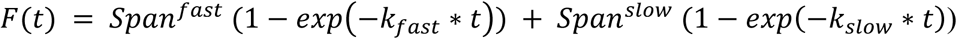

Residence time (τ) was τ = 1/k_off_, so there are two residence time, τ^fast^ and τ^slow^ corresponding to k_off_^fast^ and k ^slow^ respectively in the above equation. Bound fractions were estimated by the reduction of fluorescence signals in unbleached areas after photobleaching as soluble fraction and percentage of dynamic chromatin bound fraction was calculated by

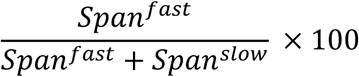

### Chromosome spread

Cells were treated with 2 μg/ml doxycycline for 24 h, followed by 500 ng/ml dTAGV-1, 100 nM 5-Ph-IAA and 100 nM JF646 for 8 h. Mitotic arrest was induced by nocodazole treatment for 4 h. To normalize STAG1^WT^ or STAG1^RA^ expression levels, cells were sorted by FACS. Immediately after sorting, cells were incubated in hypotonic buffer (0.75 M KCl) and fixed with Carnoy’s solution. Chromosome spreads were prepared on coverslips, stained with Giemsa solution (Merck) for 9 min, rinsed in tap water, air-dried, and mounted in Entellan New (Merck).

### Chromatin fractionation and immunoprecipitation

Cells synchronized in G1 or G2 were sorted by FACS to normalize the expression levels of STAG1^WT^-Halo and STAG1^RA^-Halo. Immediately after sorting, cells were lysed in extraction buffer (10 mM Tris-HCl (pH 7.5), 100 mM NaCl, 5 mM MgCl_2_, 0.2% NP40). For preparation of whole-cell extract, a portion of the lysate was treated with 0.2 U Benzonase (Merck) for 30 min at 4 °C. For chromatin fractionation, cells were centrifuged at 2,000 × *g* for 10 min, and the resulting pellet was washed five times with the same buffer. For chromatin immunoprecipitation, 10 μl of protein A magnetic beads (Veritas) were pre-coated with 2.5 μg of anti-Smc3 antibody or rabbit IgG control. Chromatin pellet was treated with 0.5 U Benzonase for 30 min at 4 °C, and then incubated with the antibody-coated beads for 2 h at 4 °C. Beads were washed three times with wash buffer (10 mM Tris-HCl, pH 7.5, 150 mM NaCl, 0.05% Tween-20), resuspended in Laemmli sample buffer, boiled, and the supernatants were analyzed by western blotting.

### Micro-C sample preparation

Cells were synchronized in G1 phase as described above and treated with 2 μg/ml doxycycline for 24 h, 500 nM dTAGV-1 and 100 nM 5-ph-IAA for 8h. STAG1^WT^- or STAG1^RA^-Halo positive cells were sorted by FACS and immediately processed for Micro-C sample preparation. For crosslinking, sorted cells were fixed in freshly prepared 1% formaldehyde (Polysciences) at room temperature for 10 min. The reaction was quenched with 2.5 M glycine for 5 min at room temperature. Cells were washed twice with PBS and subjected to a second crosslinking step with 0.3 M disuccinimidyl glutarate (DSG; Thermo Fisher) at room temperature for 45 min, followed by quenching with 2.5 M glycine for 5 min.

For MNase treatment, cell pellet was resuspended in Micro-C buffer#1 (10 mM Tris-HCl (pH 7.5), 50 mM NaCl, 5 mM MgCl_2_, 1 mM CaCl_2,_ 0.2% NP40, 1x cOmplete EDTA-free (Roche)) and incubated on ice for 20 min. After centrifugation at 5,000 × *g* for 5 min at 4 °C, chromatin was digested with 3,000 gel units of micrococcal nuclease (MNase; New England Biolabs) at 37 °C for 10 min. The reaction was stopped by centrifugation at 5,000 × *g* for 5 min at 4 °C, and the pellet was washed with ice-cold Micro-C buffer #2 (10 mM Tris-HCl (pH 7.5), 50 mM NaCl, 10 mM MgCl_2_).

For end repair, chromatin was resuspended in Micro-C end-repair mix (10 mM Tris-HCl (pH 7.5), 50 mM NaCl, 10 mM MgCl_2_, 100 µg/ml BSA, 4 mM ATP, 5 mM DTT) supplemented with 40 U of DNA Polymerase I, Large (Klenow) Fragment (New England Biolabs) and 200 U of T4 polynucleotide kinase (New England Biolabs) and incubated at 37°C for 1 h. For biotin labeling, 66 μM each of biotin-dATP (Jena Bioscience), biotin-dCTP (Jena Bioscience), dGTP (Promega), and dTTP (Promega) were added and incubate 25°C for 1 h. The reaction was stopped by adding 30 mM EDTA and incubating 65°C for 20 min.

For proximal nucleosome ligation, chromatin was centrifuged at 10,000 × *g* for 5 min at 4 °C and washed twice with ligation buffer (50 mM Tris-HCl, pH 7.5, 10 mM MgCl₂, 1 mM ATP, 10 mM DTT). Chromatin was then resuspended in ligation buffer containing 2,000 U T4 DNA ligase (New England Biolabs) and incubated at room temperature for 3 h. After ligation, chromatin was centrifuged at 10,000 × *g* for 5 min at 4 °C, resuspended in NEBuffer 1 (New England Biolabs) supplemented with 200 U exonuclease III (New England Biolabs), and incubated at 37 °C for 15 min to remove biotin-dNTPs on un-ligated ends. Then 750 µg of ProteinaseK (Roche) and 0.14% SDS were added and incubated at 55 °C for 2 h followed by incubation at 65 °C overnight for reverse crosslinking. DNA was purified by QIAquick PCR purification kit (QIAGEN).

For biotin pull-down, 5 μl Dynabeads^TM^ MyOne^TM^ Streptavidin C1 beads (Invitrogen) were washed twice with 300 μl Tween washing buffer (5 mM Tris-HCl, pH 7.5, 0.5 mM EDTA, 1 M NaCl, 0.05 % Tween20). Purified DNA was mixed with beads resuspended in 300 μl of binding buffer (5 mM Tris-HCl, pH 7.5, 0.5 mM EDTA, 1 M NaCl) and incubated for 20 min at room temperature. Beads were washed twice with Tween washing buffer. Library preparation was performed on bead-bound DNA using the NEBNext® Ultra™ II DNA Library Prep Kit for Illumina® (New England Biolabs) according to manufacturer’s protocol. After adaptor ligation, beads were washed twice with Tween washing buffer. PCR amplification was performed using NEBNext® Multiplex Oligos for Illumina® (New England Biolabs). PCR products were purified using AMPure XP beads (Beckman Coulter).

### Micro-C data analysis

Micro-C samples were processed using a standardized computational pipeline implemented in Snakemake^63^. Raw sequencing data were trimmed for adapters using Cutadapt (v3.4) with a minimum read length threshold of 50 bp. Trimmed reads were aligned to the human reference genome (hg38) using BWA-MEM (v0.7.17) with specific parameters (-5SP -T0)^64^. Raw alignments were processed using the pairtools suite (v0.3.0) to identify and filter chromatin contact pairs. The pairtools “parse” module was applied with stringent quality filters: a minimum mapping quality (MAPQ) of 40, a maximum inter-alignment gap of 30 bp, and the “5-unique walks” policy to handle multi-mapping reads. Parsed contact pairs were sorted by genomic coordinates using pairtools sort, followed by deduplication with pairtools dedup to remove PCR and optical duplicates. Valid contact pairs were extracted using pairtools split, retaining only high-quality, non-duplicate interactions for downstream analysis (Supplementary Table. 1). Finally, high-quality contact pairs were converted into contact matrices using two complementary formats: HiC format matrices were generated using Juicer Tools (v1.22.01) and, additionally, these matrices were converted to the cooler format (.mcool) using hicConvertFormat utility of HiCExplorer. Knight-Ruiz (KR) matrix balancing normalization was applied to correct for experimental biases. The Observed/Expected matrices metaplots were created using the tool coolpup.py^65^ and KR normalization. Chromatin loops were identified using the HiCCUPS algorithm (part of Juicer Tools v1.22.01) at 25-kb resolution on KR normalized contact matrices with the “--ignore-sparsity” flag to enable loop detection in regions with lower contact density. Loops greater than 4 Mb were filtered out from the analysis. Contact probability as a function of genomic distance (Fig. 7b) was calculated as an intra-chromosomal contact frequency distribution, using logarithmically increasing genomic distance bins. The insulation score (Fig. 7d) was computed using a custom Python script following the methodology described in Crane et al., 2015^66^, extracting 10-kb resolution raw matrices and using a sliding window of 100 kb × 100 kb. Finally, A/B compartment analysis was performed at 25-kb resolution using the cooltools package (v0.5.1). Compartments were identified through principal component analysis of the normalized contact matrices using the “eigs_cis” function and data from Rao et al., 2017^49^. For each chromosome, the first five eigenvectors were calculated and checked for the best correlation with GC content and H3K27ac/H3K9me3 signals (GSE96299 and GSE95968, respectively). The compartmentalization strength was defined by the formula (AA + BB) / AB, which measures intra-compartment contacts versus inter-compartment contacts.

## Acknowledgements

We are grateful to Toyonori Sakata and Katsuhiko Shirahige for their guidance on the sample preparation methods for the Micro-C experiments and for providing plasmids for lentivirus-induced expression of OSTIR1^F74G^. We thank Yuhei Goto and Kazuhiro Aoki for their technical support with microscopy in iFRAP experiments. We also acknowledge Eri Tahara, Mai Shimoura, and Shinri Aoki for their assistance with cell line establishment, protein purification, and single-molecule experiments; Dácil Alonso for preparing STAG1^RA^mutant; Tomoyuki Hara for help with lentiviral experiments; and Yoshimi Kinoshita and Makoto Yoshida and Ana Cuadrado for valuable discussions. This work was supported by JSPS Grants-in-Aid for Scientific Research/KAKENHI (KAKENHI 20H05937, 21H04767, 25H02555) and Takeda Science Foundation (2025027792) to T.N., grant PID2022-139333NB-I00 funded by MCIN/AEI/10.13039/501100011033 and by the European Regional Development Fund (ERDF-EU) to A.L., and Grant-in-Aid for JSPS Fellows (#23KJ1386) to R.S.

## Author contributions

R.S. and T.N. designed the experiments and interpreted the data. R.S. performed the experiments. D.G. and A.L. analyzed and interpreted the Micro-C data. R.S., D.G., A.L. and T.N. wrote the manuscript.

